# Emergence of Erythromycin Resistant Invasive Group A *Streptococcus* in West Virginia, United States

**DOI:** 10.1101/2022.08.08.503263

**Authors:** Lillie Powell, Soo Jeon Choi, Chloe Chipman, Megan Grund, P. Rocco LaSala, Slawomir Lukomski

## Abstract

Combination therapy with penicillin and clindamycin has been a mainstay for treatment of invasive group A *Streptococcus* (iGAS) infections yet increasing macrolide resistance may limit such treatment for strains displaying MLS_B_ phenotypes. The CDC recently reported erythromycin and clindamycin resistance rates for iGAS exceeding 20% in 2017. Here, we investigated 76 iGAS isolates from 66 patients identified at J.W. Ruby Memorial Hospital in West Virginia from 2020-2021. *emm* typing was performed using the CDC protocol and database. Resistance genes were detected by PCR and sequencing, whereas antimicrobial susceptibility testing was performed in clinical and research laboratories with standard techniques. Median patient age was 42 years (23-86 range). 76% (n=50) of isolates were simultaneously resistant to erythromycin and clindamycin, which included both inducible (n=40) and constitutive (n=9) resistance. All *emm92* (n=35) and *emm11* (n=8) isolates were erythromycin resistant, while the remaining 11% (n=7) of resistant isolates comprised 5 *emm* types. Susceptible isolates primarily included *emm89* (n=6) iGAS. Macrolide resistance was conferred by the plasmid-borne *ermT* gene in all *emm92* isolates and by chromosomally-encoded *ermA* (n=7), *ermB* (n=7), and *mefA* (n=1) in other *emm* types. Macrolide-resistant iGAS were typically resistant to tetracycline and aminoglycoside antibiotics. Here, we characterized iGAS infections affecting non-pediatric residents across West Virginia. We showed a shift in *emm*-type distribution compared to historical and national reports, and dominance of macrolide-resistant isolates which raises concern for emerging resistance to commonly-prescribed antibiotics used in treatment of iGAS infections.

---

*Streptococcus pyogenes*, also known as group A *Streptococcus* (GAS), is a ubiquitous global pathogen that produces an array of human disease, including focal infections (e.g. pharyngitis, pyoderma, *etc*.) with or without localized suppurative complications, invasive soft tissue infections (e.g. myositis, necrotizing fasciitis, *etc*.), and systemic, oftentimes fatal infections (e.g. bacteremia, toxic shock syndrome). In addition, two post-infectious complications (e.g. glomerulonephritis and rheumatic heart disease) attributable to GAS have been well-described (1–3). While GAS remains collectively susceptible to penicillin, treatment with alternative or combination therapies such as macrolides, clindamycin, and other second-line antimicrobials is common due to patient β-lactam allergies, dosing convenience, infection severity, and/or patient acuity (4). In contrast to its predictable β-lactam susceptibility, GAS resistance to other classes of antimicrobials has been increasingly reported (5–7). In the face of ongoing dissemination of the MLS_B_ [macrolide, lincosamide, and streptogramin B] and tetracycline resistant phenotypes among GAS isolates, the Centers for Disease Control and Prevention (CDC) has labeled macrolide-resistant GAS as an emerging threat of concern (8).

As one component of its Active Bacterial Core (ABC) surveillance, the CDC’s Emerging Infections Program provides ongoing laboratory and population-based assessments of GAS infections from 10 sites in the United States. Annual reports by the program estimate the incidence of invasive GAS (iGAS) infections within the US has doubled between 2009 and 2019, with total numbers of infections increasing from approximately 11,000 cases (3.6/100,000 population) to >25,000 cases (7.6/100,000 population) (9, 10). Concomitant with this, substantial increases in the proportion of iGAS isolates resistant to erythromycin and clindamycin have been reported, with overall resistance rates climbing from <10% in 2010 to near 25% by 2017 (11).

Notably, populations at risk for such macrolide-resistant iGAS infections have included predominately non-pediatric persons aged 18-64 years, with a high incidence of those with a history of intravenous drug use (IVDU) and/or persons experiencing homelessness (11).

Based on these observations, coupled both with the observed increase in annual rates of iGAS erythromycin resistance at WVU Medicine System hospitals (37% in 2019, 54% in 2020, 87% in 2021) and with the state’s extremely high per capita rate of IVDU overdose during the past several years (12), the aim of the present study was to review clinicoepidemiology of iGAS infections within our region and to characterize specific phenotypic and genotypic antimicrobial resistance traits of the available corresponding isolates.

## RESULTS

### Patients

A total of 76 GAS isolates collected from 66 patients with invasive infections between January 2020 and June 2021 were included in the study (Fig. 1, Table S1). Median patient age was 42 years (mean 45, range 23-86) with a slight male preponderance (59%). Based on addresses as listed in the medical record, geographic distribution of all 57 in-state patients spanned 20 of the 55 West Virginia counties, with three northern counties (Harrison, Marion, and Monongalia) accounting for the highest proportion (53%) likely due to their larger populations and proximity to the main WVUMed campus in Morgantown. The majority of out of state patients were also within the WVUMed catchment area from neighboring counties of Maryland (n=4) and Pennsylvania (n=5) (Fig. 1A), although one was visiting from the Midwestern US. Nine patients had insufficient details of housing status documented in their medical record, but among the remainder, eight patients (14%) were reported by a case worker to be experiencing homelessness at the time of culture. Each of these pateints did however, have a relative or emergency contact with a WV address on file for billing, which was used in Figure 1. For 64 patients with sufficient social history documented, 39 (61%) reported recent or remote IVDU.

**Fig 1.**
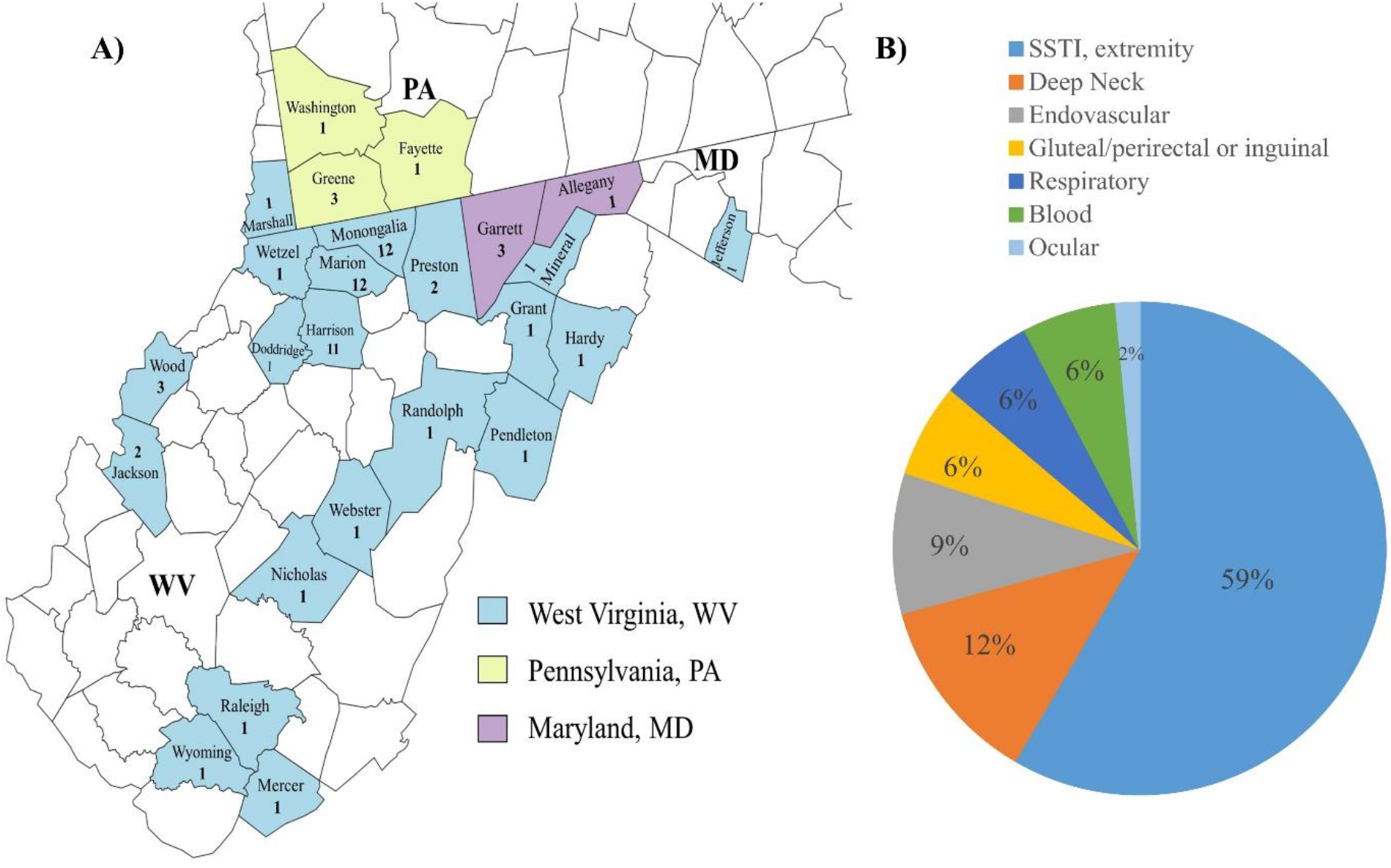
Epidemiological data of iGAS patients in this 2020-2021 collection. (A) The geographic distribution of patients. Fifty six iGAS isolates were collected from patients in 20 West Virginia (WV) counties. The residence status for 9 patients was undocumented whereas 9 patients were listed as homless. In these cases the county of residence for the billing address was used. Five isolates were from neighboring counties in Pennsylvania (PA) and 4 in Maryland (MD). (B) Assessment of infection source. The percentage rate of iGAS isolation from anatomical sources per total number of infections in the collection are shown.

Assessment of infection source for all 66 patients collectively revealed 38 patients (59%) with SSTI of the extremities, eight (12%) with infections of deep neck structures, six (9%) with endovascular sources, four (6%) with SSTI of the gluteal, perianal, sacral, or inguinal region, four (6%) with respiratory sources, one (2%) with ocular source, and four (6%) with bloodstream infections of unknown source (Fig. 1B). For one additional patient with IVDU history who presented with thrombophlebitis, bacteremia, and multiple SSTIs of the extremeties and gluteal region, the primary source could not be discerned. A total of 35 (53%) patients required surgical intervention, which included single stage debridement/washout (n=24), fasciotomy with 2-13 serial debridements (n=7), and one each with below the knee amputation revision, tricuspid valve replacement, thoracotomy with plueral decortication, and vascular thrombectomy. Eight of the remaining non-surgical patients underwent incision-drainage procedures or chest tube placement at bedside. Overall, 17 (26%) patients required at least 24hr ICU admission and at least five (7.5%) suffered mortality related to iGAS infection, though records of follow-up care were incomplete for a substantial portion of patients.

Based on review of all available culture results, iGAS infections were monomicrobial in 30 (45%) patients. By contrast, concomitant pathogens recovered from individual iGAS patients included methicillin resistant *Staphylococcus aureus* (n=19, 29%), methicillin susceptible *S. aureus* (n=8, 12%), staphylococci other than *S. aureus* (n=4, 6%), and one each harboring *S. agalactiae* and *S. pneumoniae*. An additional three patients had multiple aerobic, anaerobic, and/or facultative organisms recovered. Antimicrobial therapy varied considerably by patient acuity, duration of hospitializaiton, i.v. catheter availability, and degree of initial treatment response. A total of 51 patients (77%) received at least one dose of i.v. vancomycin or daptomycin during hospitalization, while β-lactams (amoxicillin-clavulanate, cephems, and/or penicillin) or clindamycin were used exclusively for 10 patients (15%). Another three patients who were not admitted to the hospital received no antimicrobial therapy. In total, the acuity of infection for nine patients did not require hospital admission. Another 10 patients left the hospital against medical advice prior to admission or treatment completion, nine of whom had histories of IVDU (and one who had insufficient documentation to determine IVDU history). All of these patients received prescriptions for oral clindamycin or amoxicillin-clavulanate at discharge. Among the remaining 47 patients, median hospital length of stay was 7 days (mean 14, range 1-60).

### emm types

Although surveillance across the U.S. by the Active Bacterial Core (ABC) surveilance shows an increase in iGAS disease, categorization of iGAS isolates in West Virginia was lacking. The M type of each isolate was determined by Sanger sequencing of the 5’-end of the *emm*-gene PCR product (primers in Table S2), as described (13). Analysis showed the collection was predominated by isolates of one *emm*-type; of the 66 unique patient isolates, 35 (53.0%) were *emm92* followed by *emm*-types *emm11* (n=8, 12.1%) and *emm89* (n=5, 7.7%). (Fig. 2A, Table 1). Temporal analysis of isolate *emm*-type recovery by 3-month periods showed an overall increase in isolates during April-June both years, with a significant proportion of the isolates from all quarters being *emm*-type 92 (Fig. 2A). While the presence of *emm11* and *emm89* isolates was relatively stable over time, that of *emm92* trended upward. Interestingly, the collection contained only two *emm1*, one each of *emm12* and *emm28*, and no *emm3* isolates, which historically have been correlated with a high incidence of iGAS infections (Fig. 2A) (1, 14, 15). While the data from this one-and-a-half-year study period in northern WV is not as robust as ABC-CDC national data, the findings do certainly corroborate a continual shift in *emm*-types responsible for iGAS disease, particularly among IVDU and homeless populations (11, 16).

**Table 1.**
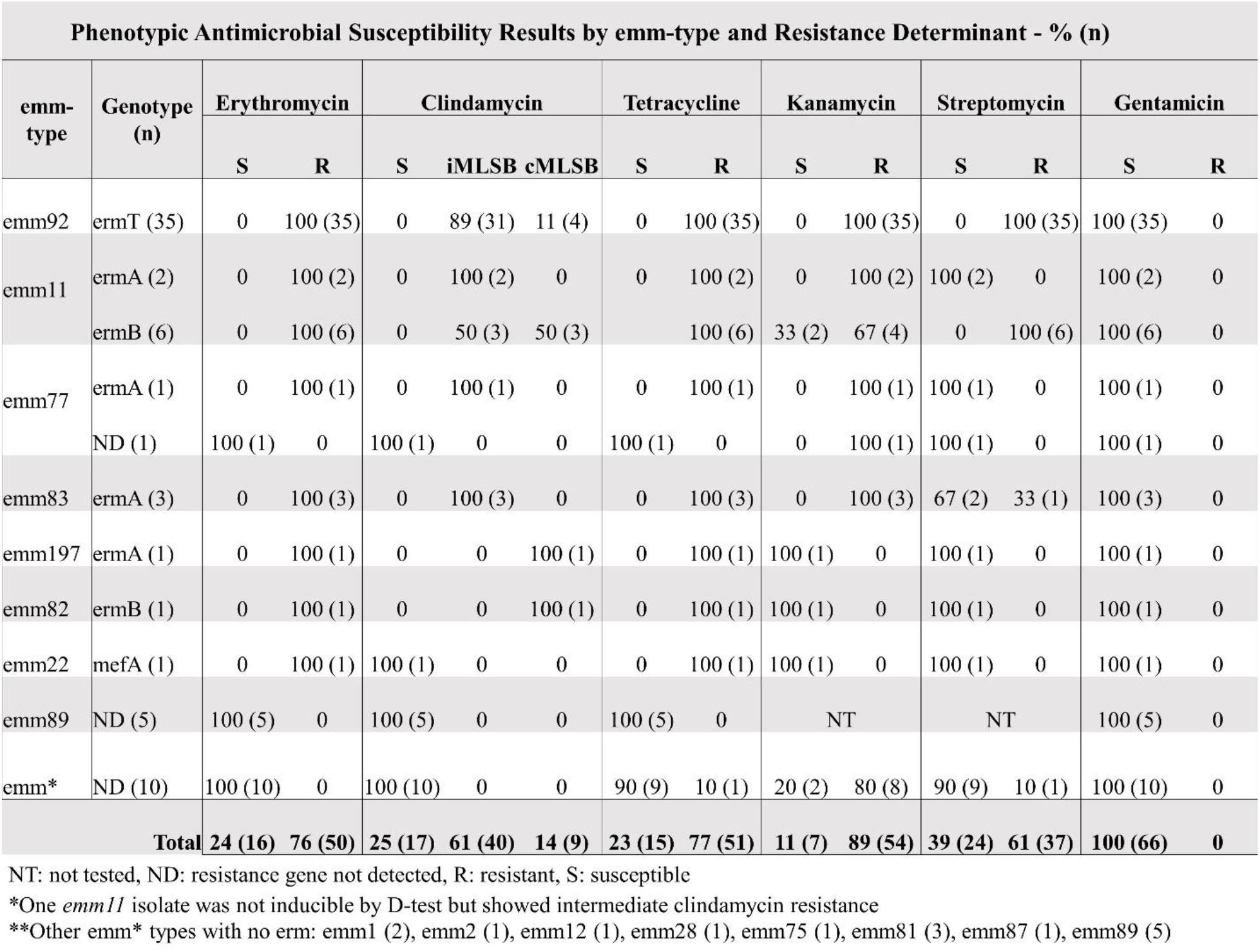
Antibiotic Resistance in invasive group A *Streptococcus* in WV

**Fig 2.**
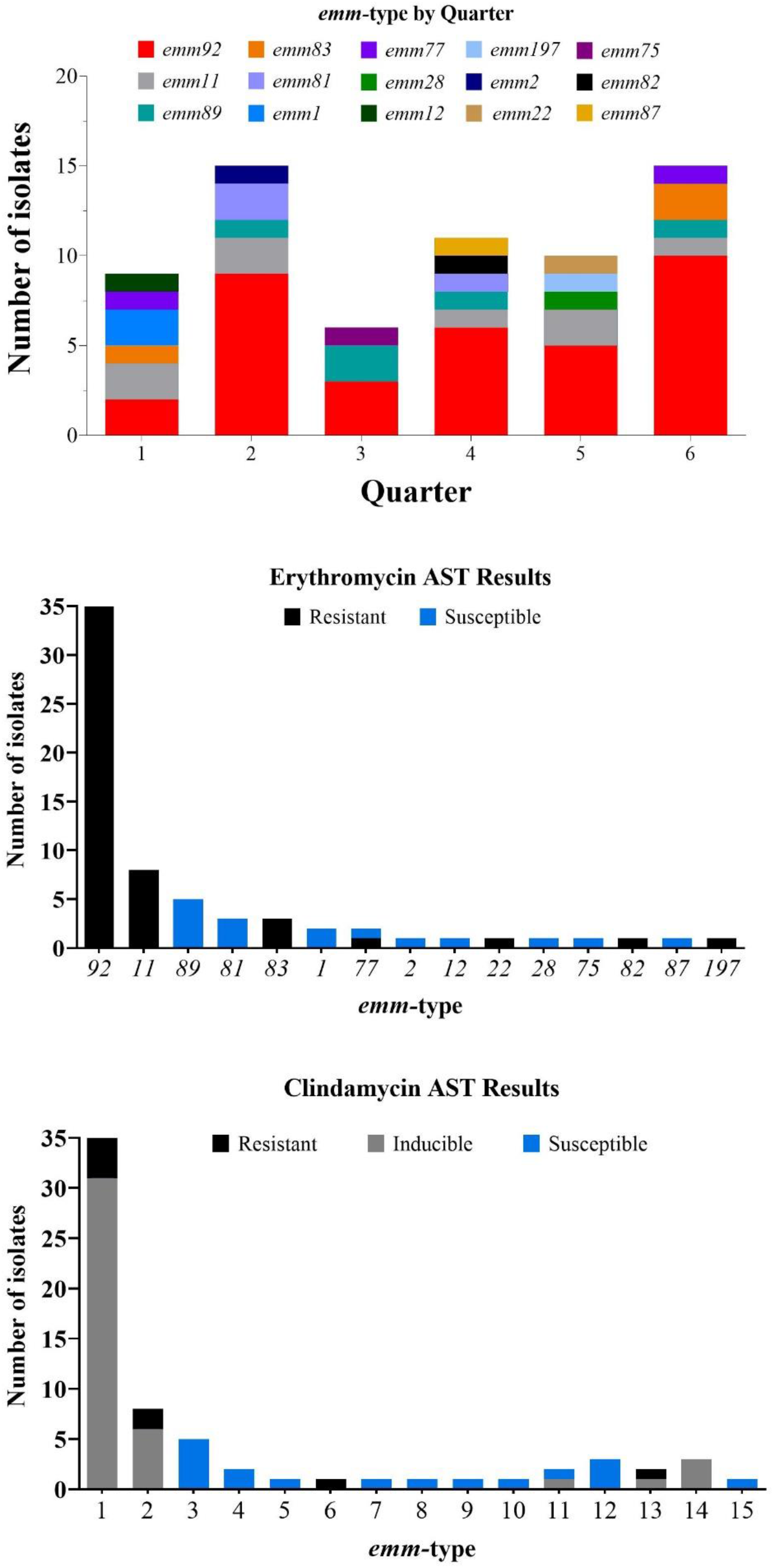
*emm-*type distribution and MLS_B_ resistance among iGAS isolates. (A) Temporal analysis of isolate *emm*-type by 3-month periods. Isolates were collected between January 2020-June 2021, which is represented by six quarters. A trend of *emm92* isolates predominating each quarter over this time period is evident. (B, C) MLS_B_ susceptibility and resistance profiles. The number of isolates resistant to erythromycin (B) and clindamycin (C) by *emm*-type was determined based on antimicrobial susceptibility testing (AST). Isolates where deemed non-susceptible to clindamycin if they had either an inducible or constitutive resistance phenotype, and susceptible in the absence of growth as determined by D-test.

### MLS_B_ susceptibility and resistance profiles

Next, we assessed antimicrobial resistance among isolates in our iGAS collection. In aggregate, 76% (50/66) of isolates were resistant to erythromycin (Table 1), which is considerably higher than reported in a larger collection (11). Aside from *emm77*, which included one erythromycin resistant and one erythromycin susceptibile isolate each, all other *emm*-types exclusively harbored either erythromycin resistant or susceptible isolates (Fig. 2B, Table 1). Disc diffusion and D-testing were used to assess clindamycin susceptibility and to determine whether non-susceptibility was constitutive or inducible (Fig. 2C). Clindamycin suceptibility mirrored that of erythromycin, with 16 erythromycin-susceptible isolates also demonstrating clindamycin susceptibility. Of the 50 erythromycin resistant isolates, 40 exhibited inducible clindamycin resistance (i.e. not detectable without erythromycin induction), nine demonstrated constitutive clindamycin resistance, and one isolate (*emm*22) was clindamycin susceptible without evidence of inhibition zone flattening. Similar to erythromycin, most *emm-* types were uniformly susceptible or resistant to clindamycin except for two *emm*77 (with one susceptible and one inducibly-resistant isolate) (Fig 2C, Table 1). Phenotypic heterogeneity was also noted among *emm*92 isolates, with four exhibiting constitutive and 31 exhibiting inducible clindamycin resistance, as well as among *emm*11 isolates with five isolates producing indicubile and three producing constitutive phenotypes (Table 1). As with *emm*-types, all replicate isolates from single patients demonstrated consistent results.

### Detection of erythromycin-resistance determinants

We next tested isolates for common erythromycin resistance genes by PCR amplification to detect the presence of the methyl transferase genes *ermA, ermB, and ermT*, as well as *mefA*, a gene associated with an efflux pump (primers in Table S2). Based on prior evidence that *ermT* is carried on the pRW35 plasmid (16), plasmid DNA was isolated from resistant isolates of various *emm*-types, yet only *emm92* isolates harbored plasmid DNA (Fig. 3A). Restriction digestion targeting a conserved *Swa*I site confirmed a ∼4.9-kb size of this pRW35-like plasmid (data not shown). All 35 *emm92* isolates were resistant to erythromycin (Table 1) and contained *ermT* detected by PCR (Fig. 3B).

**Fig 3.**
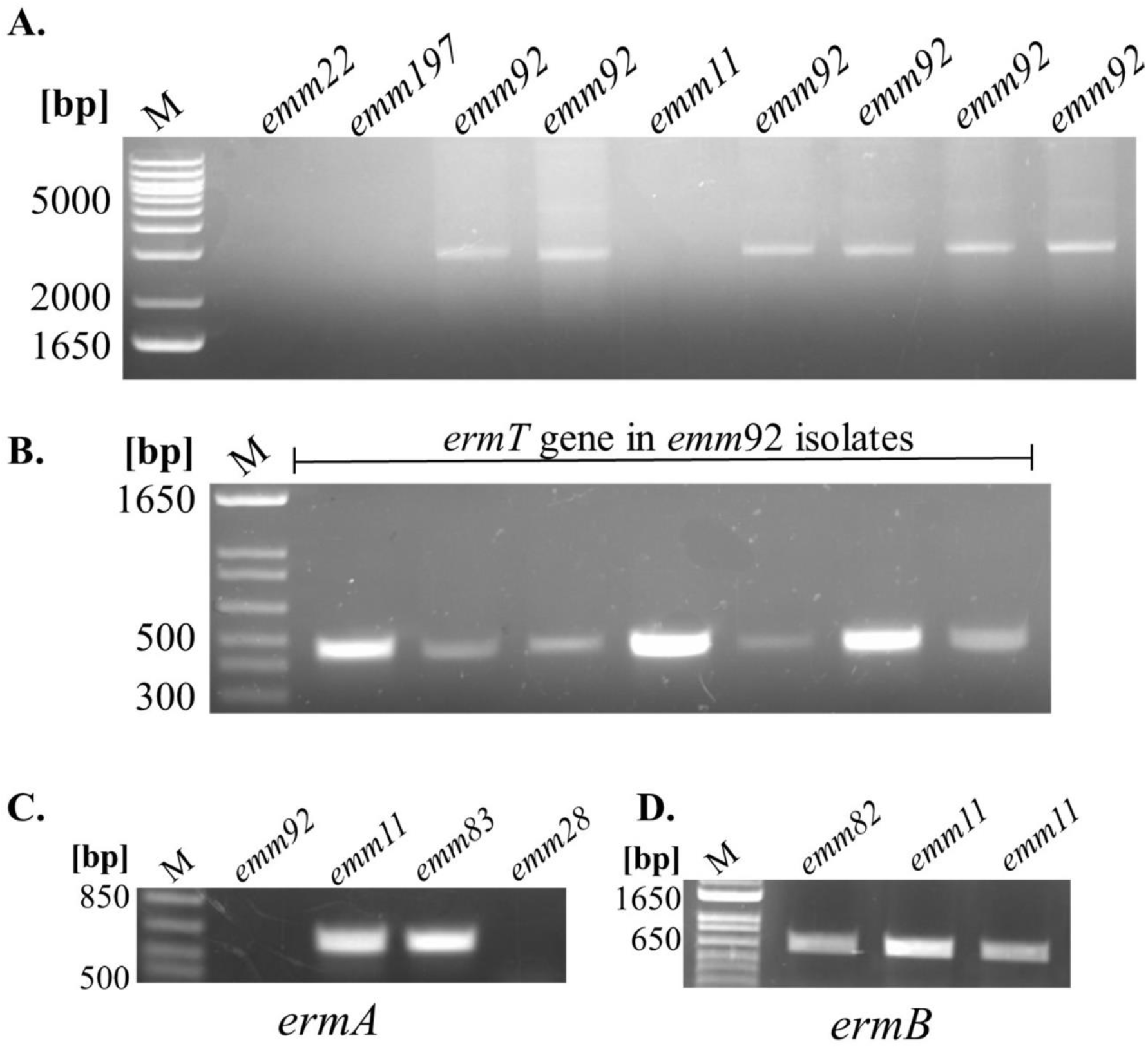
Detection of the methyl transferase genes *ermA, ermB, and ermT*. (A) Distribution of the pRW35-like plasmid among iGAS isolates. Presence of pRW35-like plasmid DNA was detected only in iGAS *emm92*-type isolates (representative samples are shown). (B) PCR detection of the *ermT*-gene. The *ermT*-specific amplicon of 452 bp was detected in *emm92* isolates using plasmid DNA as a template. (C, D) Detection of the *ermA* and *ermB* genes. Chromosomal DNA was used as a template to detect the 612-bp-*ermA* (C) and 663-bp-*ermB* (D) amplicons present in several different *emm* types. A 347-bp-*mefA* amplicon was detected in a single *emm22* isolate (not shown).

Chromosomal DNA was used for the detection of other *erm/mef* genes (Fig 3C, D). Erythromycin resistance genes identified in *emm11* isolates varied, with 6 carrying *ermB* and 2 harboring *ermA* (Table 1). Of the remaining 7 erythromycin resistant isolates of various *emm*-types, *ermA* was detected in five, *ermB* in one, and *mefA* in one. Collectively, 86% of *ermA-* containing isolates (6 of 7) showed inducible clindamycin resistance, whereas isolates containing *ermB* had a more evenly split phenotype for clindamycin resistance (3 iMLS_B_ vs 4 cMLS_B_) (Table 1). The *mefA* gene, which encodes a component of the *mefA*-*msrD* efflux pump, was detected in a single *emm*22 isolate and corresponded to the expected erythromycin resistant, clindamycin susceptible phenotype refered to as the M phenotype (17).

### Additional susceptibility and proposed resistance determinants

Isolates also underwent susceptibility testing for tetracycline by disc diffusion, as well as the aminoglycosides gentamicin, streptomycin, and kanamycin by agar dilution method using concentration ranges selected based on prior reports (5). In aggregate, 76% isolates were resistant to both tetracycline and erythromycin, whereas the single *emm87* isolate was erythromycin-sensitive but tetracycline-resistant. For aminoglycosides, all 66 isolates were susceptible to gentamicin, whereas resistance to kanamycin and streptomycin was observed in 89% and 61% of tested isolates, respectively. Importantly, in addition to their universal plasmid-encoded MLS_B_ phenotype, all *emm92* strains in this collection were uniformly resistant to tetracycline, kanamycin, and streptomycin (MICs >500 μg/mL in latter), presumably encoded by the ICESpyM92 mobile element (Table 2) (5). The remaining isolates of various *emm*-types with MLS_B_ phenotype showed resistance to either kanamycin or tetracycline.

**Table 2.** Proposed aminoglycoside and tetracycline resistance determinants

| *emm-type* | KAN<br>Resistance | STR<br>Resistance | TET<br>Resistance | Determinant | Resistance Element | Reference |
| --- | --- | --- | --- | --- | --- | --- |
| *emm92* | + | + | + | ICESpyM92, Tn916 | *Tet(M), ant(6)-Ia, aph-(3')-III* | (Sansón, Macías et al. 2019) |
|  |  |  |  | pRW35 | *ermT* | (Woodbury, Klammer et al. 2008) |
| *emm11* | + | - | + | Tn6003-like | *Tet(M), ermB, aph(3')-III* | (Sansón, Macías et al. 2019) |
|  | - | - | + | Tn6002 | *Tet(M), ermB* | (Sansón, Macías et al. 2019) |
|  | + | - | + | ICESp2905 | *Tet(O), ermA, aph-(3')-III* | (Giovanetti, Brenciani et al. 2012) |
| *emm197* | - | - | + | ICESp2905 | *Tet(O), ermA* | (Giovanetti, Brenciani et al. 2012) |
| *emm22* | - | - | + | Tn1207.1-like | *Tet(O), mefA,* | (Santagati, Iannelli et al. 2003), (Blackman Northwood, Del Grosso et al. 2009) |

## DISCUSSION

To our knowledge, the present study represents the first characterization of invasive group A *Streptococcus* (iGAS) isolates from West Virginia based on the relationship between *emm* type and macrolide resistance. The results confirm a very high rate of erythromycin resistance (76%) across seven different *emm* types producing invasive infections, the large majority of which displayed MLS_B_ phenotype and were concomitantly resistant to clindamycin. Second, a relationship between *emm* type and erythromycin resistance mechanisms in this collection became apparent. We observed that types *emm92* and *emm11* represented 53% and 12% of patient isolates, respectively, and that they accounted for 86% of the erythromycin resistant strains collectively. The third most prevalent *emm* type was *emm89*, although all isolates were susceptible to erythromycin. These findings corroborate data from the CDC’s 2010-2017 ABC nationwide surveillance program, which showed an increasing incidence of iGAS infections caused by *emm92*-, *emm11*, and *emm89*-type GAS affecting the adult U.S. population (11, 18–20). By contrast, *emm92* iGAS infections have been reported only sporadically and at much lower frequencies elsewhere (for example: (21, 22)), suggesting that expansion and dissemination of this organism may thus far be limited to the United States. Nonetheless, all three *emm* types are represented in the 30-valent M-protein based vaccine, signifying their recognition in GAS disease (23).

All the *emm92* strains in this study were found to harbor the pRW35-like plasmid containing the *ermT* gene which confers resistance to erythromycin. Plasmids harboring the highly-conserved *ermT* gene have been found in human bacteria (*S. pyogenes, S. agalactiae* (group B *Streptococcus), Lactobacillus reuteri*, and methicillin-resistant strains of *Staphylococcus aureus*) and livestock bacteria (*S. gallolyticus subsp. pasteurianus* and *Streptococcus suis*), which suggests horizontal gene transfer (16, 24–26). The practice of using animal feed containing tylosin, a macrolide additive, has been suggested as a contributing factor in the spread of *ermT* in human pathogenic organisms across different species (26). The second most commonly identified *emm* type was *emm11*, in which resistance was facilitated by either the *ermA* or *ermB* gene. In contrast to *emm92*, infections caused by the *emm11* strains displaying MLS_B_ phenotype have been broadly reported around the world (4, 27–30), including fluoroquinolone-resistant isolates in China (31). We observed 5 iGAS infections related to *emm89*. Interestingly, new acapsular *emm89* strains have emerged over the last decade as an important cause of iGAS infections worldwide (30, 32–34). Lack of hyaluronic acid capsule in those strains was compensated by the high expression of cytotoxins NADase and streptolysin O (32, 35). Finally, a single isolate, *emm22*, was found to contain the *mefA* gene as a mode of erythromycin resistance.

Antibiotic resistance to tetracycline, kanamycin, streptomycin, and gentamicin was also tested. No strains were resistant to gentamicin. However, all *emm92* strains showed resistance to kanamycin and streptomycin, as well as to tetracycline. In their previous work (5), Sanson and colleagues used whole genome sequencing to demonstrate that *emm92* isolates contain the ICESpyM92 element carrying the *Tet(M), ant(6)-Ia*, and *aph-(3’)-III* genes - which explains the resistance to tetracycline and the two aminoglycosides observed in this study – which was recently linked to increased virulence of *emm92* strains (36). Further, they showed that *emm11* isolates harboring *ermB*, and exhibiting resistance to tetracycline and kanamycin contained a Tn6003-like transposon carrying aph(3’)-III gene, while those harboring *ermB* and tetracycline resistance, only, carried Tn6002 (5). All six *emm11* isolates with *ermB* from the present work displayed one of these two phenotypes. Other studies have reported ICESp2905 as the cause of resistance to tetracycline, erythromycin, and kanamycin in *emm11* isolates containing the *ermA* gene (37, 38), which refelects the phenotypic pattern of two such isolates observed in this study collection. Our collection contained one *emm22* isolate displaying M phenotype encoded by *mefA*, which could be carried on the transposon Tn1207.3 (5, 39). Overall these results suggest that many iGAS strains in WV, especially *emm92* strains, are resistant to multiple classes of antimicrobials in addition to macrolides.

Studies of iGAS isolates throughout the U.S. have noted the emerging presence of *emm92* strains within collections (11, 18), which the present study certainly corroborates. We observed *emm92* as the predominant M-type in every quarter, although an overall decrease in incidence was noted in July-September of 2020. We reason that quarantine related to the SARS-CoV-2 pandemic may in part explain this decrease in overall number of cases, though periodic seasonality in disease incidence may also be a contributing factor. While 2010-2017 nationwide data identified *emm92, emm49*, and *emm82* as predominant iGAS-types among patients experiencing homelessness and who inject drugs (18), our 2020-2021 WV collection identified *emm92* >> *emm11* and *emm89* as top three iGAS types. In addition, CDC surveillance detected considerable numbers of historically-classical iGAS-*emm* types, (*e*.*g*., *emm1, emm12, emm28*, and *emm3)*, whereas our collection did not. Our results signify an important area for future investigation due to the emerging dominance of the *emm92*-type in iGAS infections across the United States.

In WV, drug abuse has become a serious epidemic with rates of overdose deaths rising from 1999 onward (40). Patient data from this WV cohort corroborates that IVDU is an important risk factor for resistant iGAS infections - with 60.6% of the afflicted patients reporting IVDU – compared to 8.7% reported by the CDC ABC program. Homelessness amongst West Virginians is also becoming an increasing issue. As of January 2020, it is estimated that 1,341 persons in the state experience homelessness on any given day (41). In this cohort, homelessness was reported by 13.6% of patients with others lacking documentation, whereas 12.1% of homeless patients had a history of IVDU. In comparison, a study encompassing the 10 ABC sites from 2010-2017 reported 5.8% being homeless and 6.1% having both risk factors (18).

Altogether, these findings indicate a particular vulnerability to iGAS infections associated with the socioeconomic status of U.S. citizens, which clearly affects the WV population (40). As a result, providing greater information and access to supplies for prevention of iGAS infections to those most at risk may help reduce the spread of resistant iGAS strains in the U.S.. Further studies of *emm92* iGAS isolates and categorization of resistance will be important to create better treatments and guidelines for how to prevent resistance.

### Limitations

- Although the collection of iGAS isolates reported here was derived from a broad geographic area of WV, it certainly did not include all invasive strains for the periods represented. Much of southern WV is beyond the WVUMed catchment area, and isolates from some WVUMed hospitals and other health systems would not have been captured. Further, because our hospital lab only banks invasive strains, we were unable to compare genotypic/phenotypic features of strains producing pharyngitis.
- Did not define resistance determinants for tetracycline and aminoglycosides.
- Did not explore reasons for variable inducible/consititutive phenotype within *emm* types harboring same *erm* determinant, though further work is ongoing.

## MATERIALS AND METHODS

### Bacterial isolate collection

The clinical microbiology laboratory at J.W. Ruby Memorial Hospital in Morgantown, West Virginia, serves as the primary reference facility for all 19 West Virginia University Medicine (WVUMed) System hospitals located throughout WV and in western MD, southwestern PA, and eastern OH. The WVUMed system serves an estimated patient population of 1.2 million. Most microbiological testing at J.W. Ruby Memorial Hospital and surrounding WVUMed outpatient clinics is performed by the Ruby clinical laboratory, as is referral antimicrobial susceptibility testing (AST) of many *Streptococcus* spp. from multiple system hospital laboratories. The clinical laboratory routinely banks invasive (primary and referred) isolates at −80°C for 1-2 years, with freezer space availability being the sole delimiter of storage duration. Non-invasive isolates, by contrast, including those recovered from pharyngitis cases, are not routinely held beyond seven days of specimen submission.

The strain collection for the present study included all viable primary and referred iGAS isolates available from the freezer bank, which spanned the period Jan 2020 – June 2021. For all isolates successfully retrieved, patient records were reviewed to capture demographic and selected clinical information following approval by Institutional Review Board (Protocol #2202533507). These data included: patient age, sex, and residence status, history of IVDU, ICU admission requirement, need for surgical intervention, antimicrobial regimen, and clinical outcome (Table S1). Two to four replicate isolates recovered serially from eight individual patients were included and tested separately as a quality control check. In all instances, intra-patient phenotypic and genotypic results were consistent. As a result, isolates are reported per patient throughout.

### Chromosomal and plasmid DNA isolation

Genomic DNA was isolated from a 10-μL loopful of bacteria grown in THY broth using the DNA extraction procedure, as described previously (42). Plasmid DNA was isolated using Gene JET Plasmid Miniprep Kit (Thermoscientific), with an additional cell-digestion step (1 mg/mL lysozyme and 0.5U/µl mutanolysin) at 37°C for 1 hour. Plasmid DNA, uncut and digested with *Swa*I, was analyzed on a 0.8% agarose gel to detect supercoiled and linear forms of plasmid pRW35-like found in *emm92* strains (16).

### Identification of resistance genes *erm/mef* and *emm* typing

Plasmid DNA was used as a PCR template with the *ermT*-specific primers whereas genomic DNA was used as the PCR template with primers detecting the *ermA(TR), ermB(AM)*, and *mefA* genes (Table S2). Control strains harboring defined resistance determinant were acquired from the Centers for Disease Control and Prevention’s (CDC) Emerging Infection Program laboratory. Genomic DNA was also used to determine isolate *emm* type by Sanger sequencing of the PCR products generated with primers *emm*1b and *emm*2 (13), followed by the BLAST search on the database at the Streptococcal Epidemiology Laboratory at the CDC.

### Antimicrobial Susceptibility Testing

Erythromycin, clindamycin, and tetracycline susceptibility testing was performed in the clinical microbiology laboratory using methods described by the Clinical Laboratory Standards Institute (CLSI M100-S31). In brief, all automated testing utilized Vitek 2 ST-02 cards (BioMeriuex, Durham, NC) and was performed historically upon isolate recovery for clinical management purposes. All valid results were reported to and retrieved from patient electronic health records (Epic, Verona, WI). All historic testing met ongoing quality control criteria as outlined in the laboratory Quality Management Plan and Individualized Quality Control Plan as required for accreditation. Subsequently, disc diffusion and D-testing were performed over multiple days using thawed isolates from the freezer bank. Following two serial propagations on sheep blood agar, swabs of 0.5 MacFarland suspensions prepared from isolated colonies were inoculated for confluent growth on cation-adjusted Mueller Hinton agar with 5% sheep blood (BBL, Becton-Dickinson, Franklin Lakes, NJ) using discs (BD) containing conventional drug masses. Quality control organisms, including ATCC BAA-977 (*Staphylococcus aureus* with inducible MLS_B_), ATCC BAA-976 (*Staphylococcus aureus* without inducible MLS_B_), and ATCC 49619 (*Streptococcus pneumonia*), were tested in parallel each day of use. Plates were incubated at 35°C in 5% CO_2_ environment for 20-24 hrs before measuring zone of inhibition with a manual caliper in reflected light. Zone diameters were interpreted using CLSI clinical breakpoints, and any degree of clindamycin zone flattening in proximity to erythromycin disc was interpreted as a positive D-test result.

Susceptibility testing against aminoglycosides (gentamicin, kanamycin, and streptomycin) was performed by agar dilution on Mueller Hinton media (Becton-Dickinson Laboratories) prepared in the research laboratory. A saline suspension of each isolate was prepared and adjusted to achieve an absorbance of 1 Klett unit and a 10-µL drop (∼10^4^ CFU) of the cell suspensions was plated in singlicate onto agar medium containing arbitrarily selected concentrations ranging from 50 to 500 μg/mL, as described (5)(Table S3). Plates were incubated at 37°C in 5% CO_2_ overnight then observed for presence or absence of growth.

## ACKNOWLEDGEMENTS

We thank the Centers for Disease Control and Prevention’s (CDC) Emerging Infection Program laboratory for providing control GAS strains. S.L. and M.E.G. acknowledge funding from a grant awarded as a result of Broad Agency Announcement (BAA) HDTRA1-14-24-FRCWMD-Research and Development Enterprise, Basic and Applied Sciences Directorate, Basic Research for Combating Weapons of Mass Destruction (C-WMD), under contract #HDTRA1035955001, and in part by the Vaccine Development Center at WVU-HSC, Research Challenge Grant no.HEPC.dsr.18.6 from the Division of Science and Research, WV Higher Education Policy Commission, and by the Transition Grant Support; Office of Research and Graduate Education, WVU Health Sciences Center (to S.L.). LP and CP are supported by the Department of Microbiology, Immunology and Cell Biology Research Internship for Undergraduates in the Immunology and Medical Microbiology degree program.

## Supplementary material

**Table S1.** Isolate collection information.

| Patient # | *emm*-Type | age (yr) | Sex | Homeless | IVDU | Source of infection | Specimen | ICU | Surgery (No.) |
| --- | --- | --- | --- | --- | --- | --- | --- | --- | --- |
| 1 | *emm1* | 71 | male | no | no | Respiratory | Blood | no | no |
| 2 | *emm1* | 86 | female | no | no | Respiratory | Sterile fluid | yes | no |
| 3 | *emm2* | 67 | male | no | no | SSTI | Blood | no | no |
| 4 | *emm11* | 46 | female | no | yes | SSTI | Extremity | no | yes |
| 5 | *emm11* | 24 | female | no | yes | Deep neck | Neck aspirate | no | no |
| 6 | *emm11* | 33 | male | no | no | SSTI | Extremity | no | yes |
| 7 | *emm11* | 41 | male | no | no | SSTI | Gluteal/perirectal or inguinal | no | yes |
| 8 | *emm11* | 59 | male | no | no | SSTI | Extremity | no | no |
| 9 | *emm11* | 50 | male | no | yes | SSTI | Extremity | no | yes |
| 10 | *emm11* | 43 | male | yes | yes | Endovascular | Blood | yes | yes |
| 11 | *emm11* | 35 | male | no | yes | SSTI | Gluteal/perirectal or inguinal | no | yes |
| 12 | *emm12* | 47 | male | no | no | unknown | Blood | no | no |
| 13 | *emm197* | 60 | male | no | no | SSTI | Gluteal/perirectal or inguinal | no | no |
| 14 | *emm22* | 30 | female | unknown | yes | Endovascular | Blood | no | no |
| 15 | *emm28* | 60 | male | no | no | SSTI | Blood | yes | no |
| 16 | *emm75* | 28 | female | no | no | Ocular | Sterile fluid | no | no |
| 17 | *emm77* | 50 | male | yes | yes | unknown | Blood | no | no |
| 18 | *emm77* | 32 | female | no | yes | Respiratory | Sterile fluid | yes | yes |
| 19 | *emm81* | 30 | male | no | yes | Endovascular | Blood | no | no |
| 20 | *emm81* | 34 | female | yes | yes | SSTI | Extremity | yes | yes |
| 21 | *emm81* | 59 | male | yes | no | SSTI | Extremity | yes | yes (13) |
| 22 | *emm82* | 25 | male | no | yes | Endovascular | Blood | no | no |
| 23 | *emm83* | 59 | female | unknown | unknown | unknown | Blood | yes | no |
| 24 | *emm83* | 42 | female | no | yes | SSTI | Gluteal/perirectal<br>or inguinal | no | yes |
| 25 | *emm83* | 51 | male | no | yes | SSTI | Extremity | no | yes (4) |
| 26 | *emm87* | 36 | female | no | no | Deep neck | Neck aspirate | no | no |
| 28 | *emm89* | 71 | female | no | no | Respiratory | Sterile fluid | yes | yes |
| 29 | *emm89* | 66 | male | no | no | SSTI | Extremity | no | no |
| 30 | *emm89* | 46 | male | no | no | SSTI | Extremity | no | yes |
| 31 | *emm89* | 59 | male | unknown | no | SSTI | Extremity | no | no |
| 32 | *emm92* | 44 | female | unknown | yes | SSTI | Extremity | no | no |
| 34 | *emm92* | 48 | female | yes | yes | SSTI | Extremity | no | yes (4) |
| 37 | *emm92* | 62 | male | no | no | SSTI | Blood | no | yes |
| 38 | *emm92* | 63 | male | no | no | SSTI | Blood | no | no |
| 40 | *emm92* | 25 | male | no | yes | Endovascular | Blood | yes | yes |
| 41 | *emm92* | 26 | male | no | yes | SSTI | Extremity | no | no |
| 42 | *emm92* | 35 | male | unknown | yes | Deep neck | Neck aspirate | no | yes |
| 43 | *emm92* | 37 | male | no | yes | SSTI | Extremity | no | yes |
| 45 | *emm92* | 28 | male | no | yes | Deep neck | Neck aspirate | no | yes |
| 46 | *emm92* | 42 | male | no | yes | SSTI | Extremity | yes | yes |
| 47 | *emm92* | 23 | male | yes | yes | SSTI | Extremity | no | no |
| 48 | *emm92* | 78 | female | no | no | SSTI | Sterile fluid | yes | yes |
| 49 | *emm92* | 29 | male | no | yes | Endovascular | Blood | yes | yes |
| 50 | *emm92* | 23 | female | no | yes | SSTI | Extremity | no | yes |
| 51 | *emm92* | 42 | male | no | yes | SSTI | Extremity | yes | no |
| 54 | *emm92* | 51 | male | no | no | Deep neck | Neck aspirate | no | yes |
| 55 | *emm92* | 28 | female | unknown | yes | unknown | Blood | no | no |
| 56 | *emm92* | 31 | male | no | yes | SSTI | Extremity | no | yes |
| 57 | *emm92* | 23 | female | unknown | yes | SSTI | Extremity | no | no |
| 58 | *emm92* | 42 | female | yes | yes | SSTI | Extremity | no | no |
| 59 | *emm92* | 71 | female | no | no | Deep neck | Blood | yes | no |
| 60 | *emm92* | 35 | female | no | yes | SSTI | Extremity | no | yes |
| 61 | *emm92* | 83 | female | no | no | SSTI | Blood | no | no |
| 62 | *emm92* | 39 | male | no | yes | SSTI | Extremity | no | no |
| 63 | *emm92* | 23 | female | no | yes | SSTI | Extremity | no | yes |
| 64 | *emm92* | 45 | male | unknown | unknown | Deep neck | Neck aspirate | no | no |
| 65 | *emm92* | 34 | female | unknown | yes | SSTI | Extremity | no | yes |
| 66 | *emm92* | 60 | male | yes | yes | SSTI | Extremity | no | no |
| 27* | *emm89* | 55 | male | no | no | SSTI | Blood | no | no |
| 27* | *emm89* | 55 | same | same | same | same | Extremity | same | same |
| 33* | *emm92* | 33 | female | no | yes | SSTI | Blood | no | yes |
| 33* | *emm92* | same | same | same | same | same | Extremity | same | same |
| 35* | *emm92* | 47 | male | no | no | SSTI | Extremity | same | yes (3) |
| 35* | *emm92* | same | same | same | same | same | Blood | same | same |
| 36* | *emm92* | 58 | female | no | no | SSTI | Extremity | yes | yes (8) |
| 36* | *emm92* | same | same | same | same | same | Blood | same | same |
| 39* | *emm92* | 34 | female | no | yes | Deep neck | Sterile fluid | yes | yes |
| 39* | *emm92* | same | same | same | same | same | Neck aspirate | same | same |
| 44* | *emm92* | 30 | female | no | yes | SSTI | Extremity | no | yes (4) |
| 44* | *emm92* | same | same | same | same | same | Extremity | same | same |
| 52* | *emm92* | 44 | male | yes | yes | SSTI | Extremity |  | yes |
| 52* | *emm92* | 44 | same | same | same | same | same | same | same |
| 53* | *emm92* | 40 | male | no | yes | Endovascular | Blood | no | no |
| 53* | *emm92* | 40 | same | same | same | same | Blood | yes | yes |
| 53* | *emm92* | same | same | same | same | same | Gluteal/perirectal<br>or inguinal | same | same |
| 53* | *emm92* | same | same | same | same | same | Extremity | same | same |

**Table S2:**
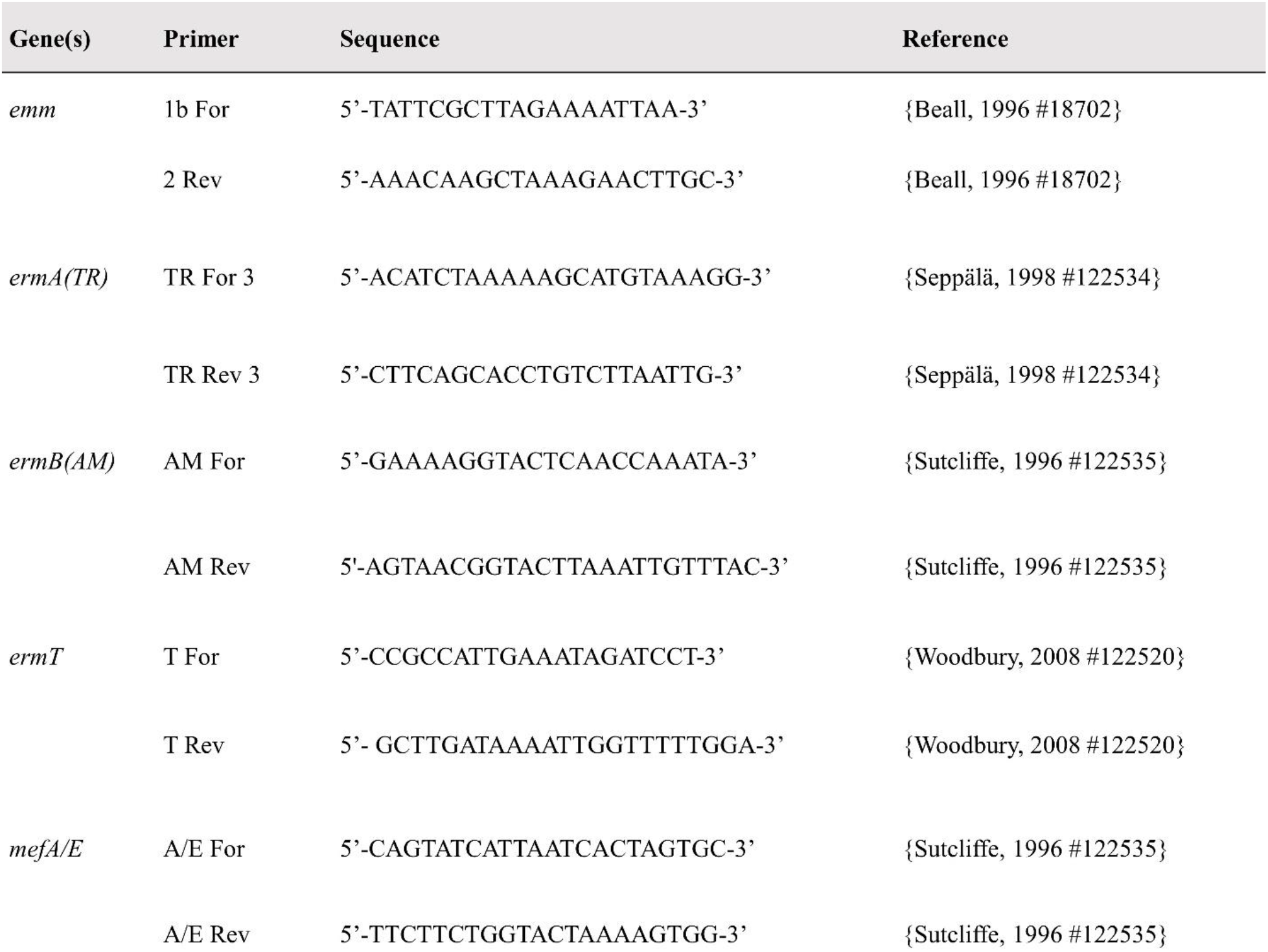
Primers used for detection of *emm*-type, erythromycin resistance gene cassettes, and regulators.

